# Characterization of the dimeric interactions of dimeric and tetrameric conformations of the PvNV protrusion-domain using a mixed DFT/QTAIM approach

**DOI:** 10.1101/2020.02.20.957316

**Authors:** Elahe K. Astani, Nai-Chi Chen, Yen-Chieh Huang, Sara Ersali, Pei-Ju Lin, Hong-Hsiang Guan, Chien-Chih Lin, Phimonphan Chuankhayan, Chun-Jung Chen

**Affiliations:** Life Science Group, Scientific Research Division, National Synchrotron Radiation Research Center, Hsinchu, Taiwan; Department of Chemistry, Faculty of Science, Tarbiat Modares University, Tehran, Iran; Babeș-Bolyai University, Faculty of Chemistry and Chemical Engineering, Cluj-Napoca, Romania; Department of Biotechnology and Bioindustry Sciences, National Cheng Kung University, Tainan City, Taiwan; Department of Physics, National Tsing Hua University, Hsinchu, Taiwan

**Keywords:** P-domain of PvNV, dimeric and tetrameric conformations, dimeric interfaces, DFT, NBO, QTAIM, hydrogen bond, hydrophobic, charge-charge, charge-dipole, dipole-dipole interactions

## Abstract

The protrusion-domain (P-domain) of *Penaeus vannamei* nodavirus (PvNV) exists as two dimer-dimer conformations: one is a protein dimer and the other is a protein tetramer. We undertook a theoretical study to gain a clear understanding of the nature of the stabilizing interactions at the dimeric interfaces of the dimeric and tetrameric conformations of the PvNV P-domain (PvNVPd) using the quantum theory of atoms in molecules (QTAIM) and natural-bond orbital (NBO) analyses in the framework of the density-functional theory (DFT) approach. The QTAIM analysis characterized the presence of multiple hydrogen bonds of common types with strength ranging from electrostatic to the covalent limit inside the PvNVPd dimer-dimer interfaces. Val257-Lys335, Phe294-Val330, Gln296-Thr328, Glu296-Thr329, Thr328-Gln297, Val330-Ala293, Lys335-Asp256 and Lys335-Val257 pairs are critical residue pairs of all three dimeric interfaces of PvNVPd. They preserve these dimeric interfaces through charge-charge, charge-dipole, dipole-dipole, hydrophobic and hydrogen bond interactions. The strongest intermolecular dimer–dimer interactions belong to the dimeric interface between subunits A and B of PvNVPd in the tetrameric conformation.

## Introduction

*Penaeus vannamei* nodavirus (PvNV) is a non-enveloped viruses that belongs to the Nodaviridae family; it can cause 100% mortality in larval, post-larval and early juvenile stages resulting in great economic loss in prawn hatcheries [1]. In the past decade, PvNV has spread to many Asian and Oceanic countries, including China, India, Taiwan, Thailand, Malaysia, Indonesia and Australia [2,3]. The Nodaviridae genome consists of two single-stranded positive-sense short-genomic RNAs encoding three gene products. RNA 1 (3.2 kb) encodes the RNA-dependent RNA polymerase for RNA replication and the nonstructural B2 protein for the host RNA interference suppressor; RNA 2 (1.2 kb) encodes the viral capsid protein (CP) for viral capsid assembly [4–6]. Previous studies exhibited that the recombinant PvNV CP of full-length 368 amino acids assembles into virus-like particles (VLPs) in the *T*=3 icosahedral capsids of diameter approximately ∼35-40 nm. The high-resolution 3D reconstructions of the *T*=3 PvNV VLPs, solved at 3.7 Å resolution, reveals that one CP comprises four regions, including the N-terminal arm (N-arm), the shell domain, the linker and the protrusion domain (P-domain) (residues 250-368) [7,8]. Crystal structures of the PvNV P-domain (PvNVPd) exist in two distinct dimer-dimer conformations of which one possesses one dimer (space group *P*2_1_) and the other has two dimers (*P*2_1_2_1_2_1_) [7].

To gain insight into the dimer-dimer interfaces of PvNV P-domains, one must characterize accurately the nature of the non-covalent intermolecular interactions involved in these interfaces using applicable computational methods. These dimeric interactions comprise hydrogen-bonding (H-bonding), electrostatic, and van der Waals interactions [9,10] that all of them fall under the umbrella of Coulombic interactions [11–13]. Among them, hydrogen bonds (H-bonds) play the most important role in maintaining the dimeric PvNVPd interfaces. Bader’s quantum theory of atoms in molecules (QTAIM) [14,15] and natural bond orbital (NBO) analysis [16,17] are two extremely useful theoretical methods to provide a clear understanding of the physical nature of the H-bonding interactions. Zhao and Truhlar have confirmed the suitability of M06 functionals of density-functional theory (DFT) in investigating the H-bonding interactions of hydrogen-bonded systems [18–20]. Our primary objective in this study is to identify accurately the nature of the dimeric interactions within the dimeric P-domain interfaces of each PvNV conformation with QTAIM and NBO analyses in the framework of a DFT approach. The other purposes are to compare the stabilities of the interfaces and to identify the most stable conformation.

## Results and discussion

The conformation of the PvNV P-domain of space group *P*2_1_2_1_2_1_ contains two A/B and C/D dimers. The crystal structure of this tetrameric protein is available in Protein Data Bank (PDB; accession code 5YL0) (Fig 1). The conformation of the PvNV P-domain of space group *P*2_1_ is a dimeric protein (PDB code 5YKZ) and has a parallel shaped model [7] (Fig 2). All three dimeric interfaces in these two proteins have the same interacting residues consisting of Asp256, Val257, Ala293, Phe294, Leu295, Glu296, Gln297, Asn298, Gln300, Pro325, Thr326, Thr328, Thr329, Val330, Ser331, Lys335, Ile364 and Ala366. These residues are located 5 Å from each other and are arranged in disparate spatial orientations with respect to each other within each dimeric interface. To identify the physical nature of the dimeric interactions, we constructed a structural model from each dimeric interface on separating its specified residues from the other parts of the corresponding protein. Fig 3 depicts the dimeric interface between subunits A and B in the tetrameric conformation of PvNVPd.

**Fig 1.**
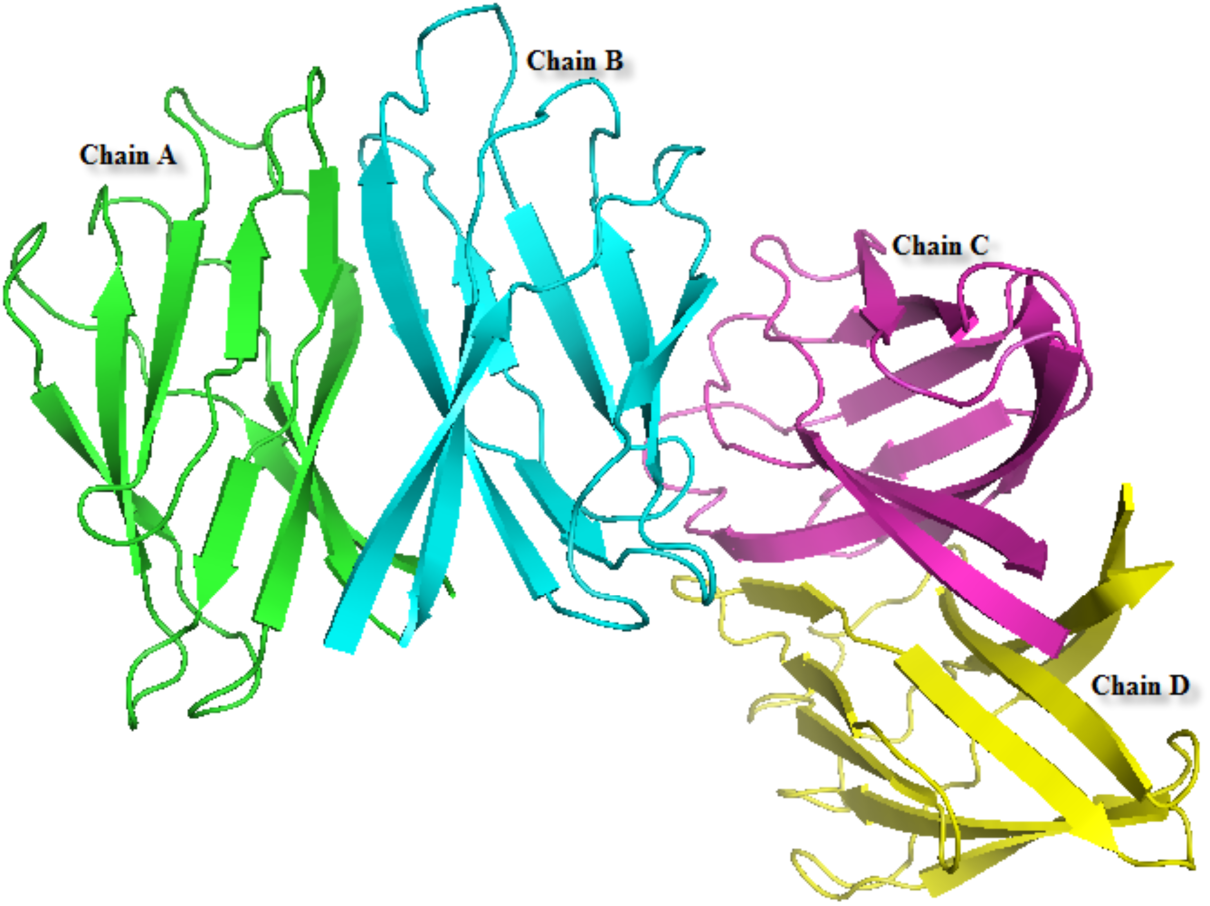
PvNV P-domain conformation with space group *P*2_1_2_1_2_1_ is a tetrameric protein containing two dimeric A/B and C/D interfaces.

**Fig 2.**
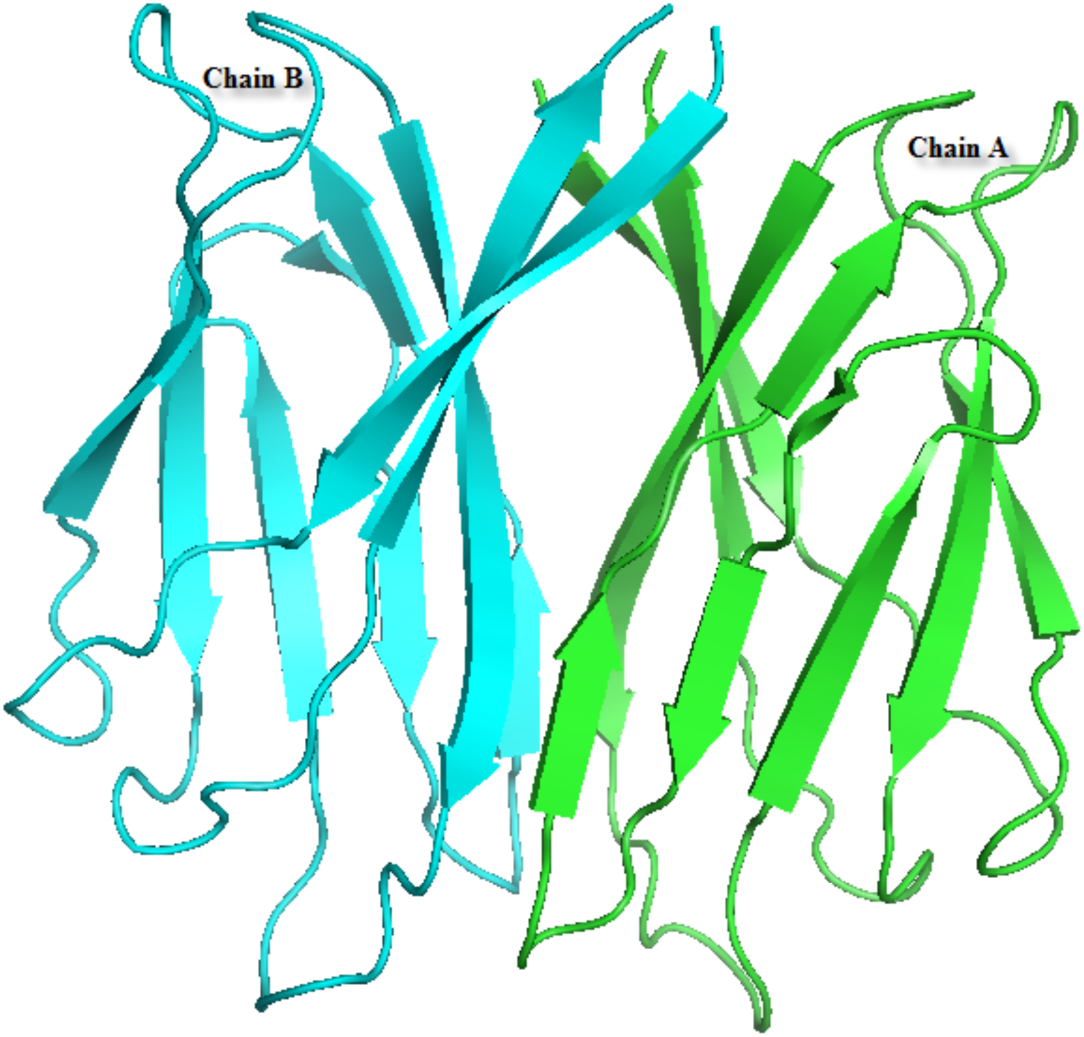
The conformation of the PvNV P-domain of space group *P*2_1_ has a dimeric A/B interface and exhibits a parallel shape.

**Fig 3.**
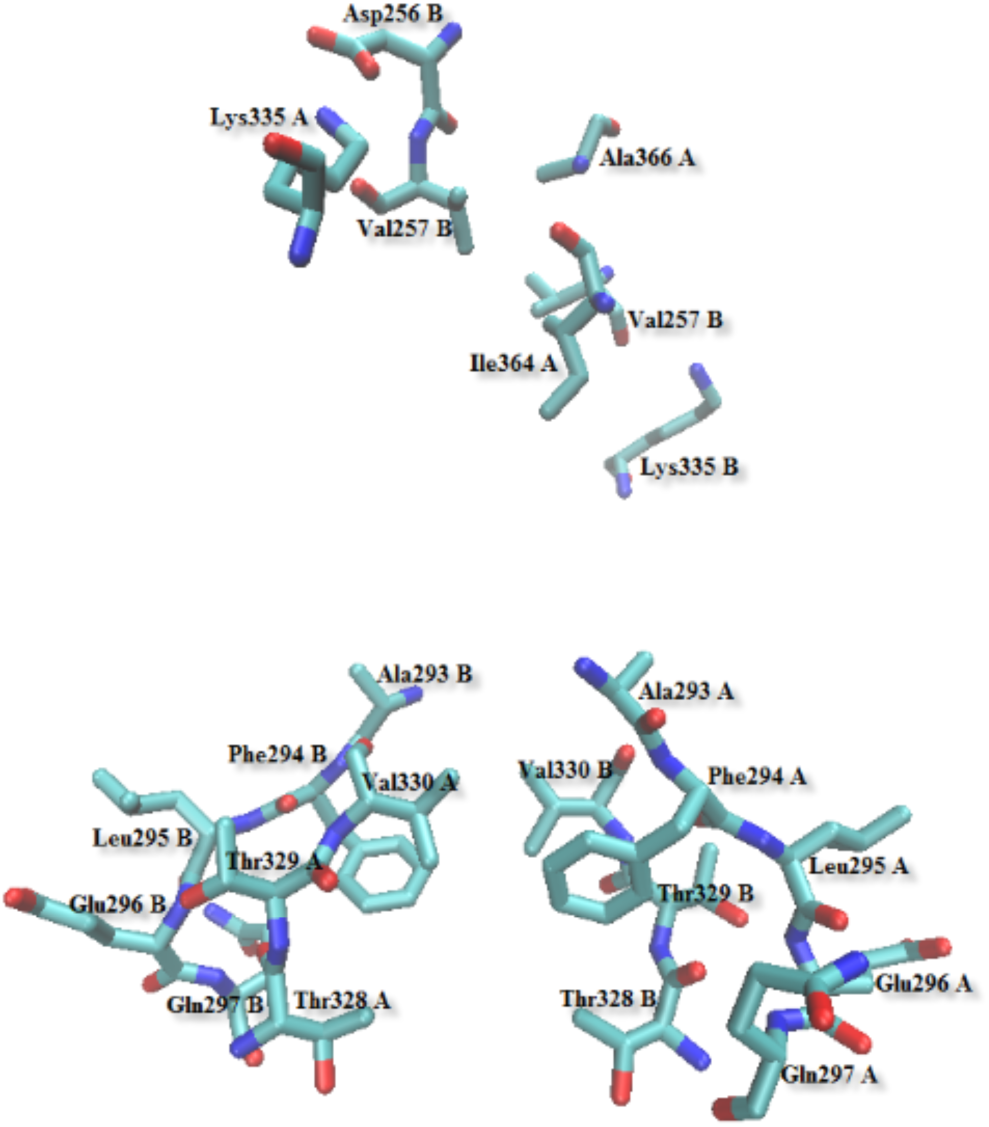
Structural model of residues involved in dimeric interactions within the dimeric A/B interface of the PvNV tetrameric P-domain.

QTAIM analysis is a reliable theoretical tool to understand better the physical nature of the intra- and the intermolecular interactions in terms of the topological features of the distribution of electron density, *ρ*(r), the bond path (BP) and bond critical point (BCP) [14,15]. The magnitude of *ρ*(r) at a BCP, *ρ*_BCP_ (**r**_cp_), its Laplacian, ∇^2^*ρ*_BCP_ (**r**_cp_), and the density of electronic energy, *H*_BCP_, provide valuable information about the nature and strength of shared (covalent bonds) or closed-shell (such as, van der Waals, ionic, H-bonding, H-H bonding, etc.) interactions [21,22].

According to NBO theory, the donor-acceptor interaction is associated with the charge transfer, CT, from a lone electron-pair orbital of a proton acceptor (electron donor), *n*_B_, to a valence antibonding orbital of a proton donor (electron acceptor), σ*_A-H_. The *n*_B_ → σ*_A-H_ orbital overlap is characteristic for the H-bonding interaction [17]. The energy of the *n*_B_ → σ*_A-H_ interaction is called the second-order stabilization energy, *E*^(2)^, that is evaluated with the second-order perturbation theory according to the below equation [16,17,23]:

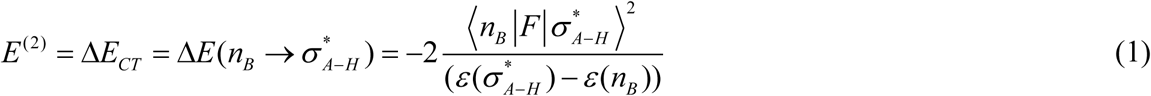

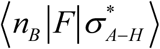 is the Fock matrix element; 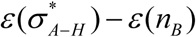 is the energy difference between the donor and the acceptor orbitals. As the stabilization energy estimates the magnitude of the strength of the donor-acceptor interaction upon the formation of a H-bond, it can serve as an energetic criterion to evaluate the strength of the H-bonding interaction.

As mentioned above, the tetrameric protein has two dimeric interfaces of which one is between subunits A and B, named model I, and the other is between subunits C and D, called model II. The dimeric interface between subunits A and B of parallel PvNVPd is termed model III. In the following sections, the dimeric interactions in each of these models is explored to compare their stabilities and to identify the most stable model by QTAIM and NBO analyses.

### QTAIM analysis on the dimeric interfaces of PvNV tetrameric P-domain

The calculated topological parameters of the QTAIM analysis of models I and II are collected in Table 1. From the topological criteria of the Koch–Popelier point of view [24], if *ρ*_BCP_ on the bond path between a hydrogen atom and a proton acceptor (H…B BP) ranges from 0.002 to 0.040 a.u. and its ∇^2^*ρ*_BCP_ lies in range 0.020-0.150 a.u., these values agree with the presence of a H-bonding interaction at the BCP. The H-bonding interaction energy (*E*_HB_) is an appropriate energetic quantity that can serve as a criterion to measure the strength of the H-bond. Espinosa and coworkers performed a topological and related energetic analysis on a range of hydrogen-bonded complexes and found a correlation between the *E*_HB_ with the electronic potential energy density at the BCP, *V*_BCP_, by the expression [25,26]:

**Table 1.** H-bond distance (*d*) and selected topological parameters of the electron density in various H…B BCPs identified within the dimeric A/B (model I) and C/D (model II) interfaces of the tetrameric protein, calculated at the M06-2X/6-31G** level.

| Model | Residue pair | H-bond | $d$<br>(Å) | $\rho_{\text{BCP}}$<br>(a.u.) | $\nabla^2\rho_{\text{BCP}}$<br>(a.u.) | $H_{\text{BCP}}$<br>(a.u.) | $\vert E_{\text{HB}}\vert $<br>(kJ/mol) |
| --- | --- | --- | --- | --- | --- | --- | --- |
| Model I | Ala293 A-Val330 B | Cα-Hα...O | 2.43 | 0.0122 | 0.0446 | -0.0080 | 11.88 |
| Model II | - | - | - | - | - | - | - |
| Model I | Phe294 A-Thr328 B | Cδ2-Hδ2...O | 2.66 | 0.0075 | 0.0269 | -0.0041 | 6.48 |
| Model II | Phe294 C-Thr328 D |  | 2.64 | 0.0069 | 0.0254 | -0.0035 | 5.88 |
| Model I | Phe294 A-Val330 B | N-H...O | 1.84 | 0.0302 | 0.1137 | -0.0240 | 33.41 |
| Model II | Phe294 C-Val330 D |  | 2.16 | 0.0167 | 0.0597 | -0.0163 | 20.82 |
| Model I | Leu295 A-Thr329 B | Cδ1-Hδ13...Oγ1 | 2.62 | 0.0082 | 0.0261 | -0.0048 | 7.08 |
| Model II | Leu295 C-Thr329 D |  | 2.53 | 0.0097 | 0.0295 | -0.0062 | 8.64 |
| Model I | Glu296 A-Thr329 B | N-H...Oγ1 | 1.80 | 0.0358 | 0.1268 | -0.0287 | 38.98 |
| Model II | Glu296 C-Thr329 D |  | 1.89 | 0.0280 | 0.0915 | -0.0233 | 30.43 |
| Model I | Gln297 A-Thr328 B | N-H...O | 1.74 | 0.0328 | 0.1145 | -0.0253 | 34.67 |
| Model II | Gln297 C-Thr328 D |  | 1.88 | 0.0277 | 0.0927 | -0.0219 | 29.28 |
| Model I | - | - | - | - | - | - | - |
| Model II | Pro325 C-Gln300 D | Cβ-Hβ2...Oε1 | 2.67 | 0.0064 | 0.0221 | -0.0033 | 5.27 |
| Model I | - | - | - | - | - | - | - |
| Model II | Thr326 C-Asn298 D | Cα-Hα...Oδ1 | 2.55 | 0.0078 | 0.0271 | -0.0044 | 6.84 |
| Model I | Thr328 A-Gln297 B | N-H...O | 1.72 | 0.0429 | 0.1648 | -0.0360 | 49.52 |
| Model II | Thr328 C-Gln297 D |  | 1.87 | 0.0293 | 0.1004 | -0.0241 | 32.04 |
| Model I | Thr328 A-Gln297 B | Oγ1-Hγ1...O | 2.39 | 0.0085 | 0.0337 | -0.0060 | 8.96 |
| Model II | - | - | - | - | - | - | - |
| Model I | - | - | - | - | - | - | - |
| Model II | Val330 C-Phe294 D | N-H...O | 1.95 | 0.0227 | 0.0778 | -0.0184 | 24.58 |
| Model I | Lys335 A-Asp256 B | Nζ-Hζ1...Oδ1 | 1.61 | 0.0588 | 0.1466 | -0.0557 | 64.81 |
| Model II | - | - | - | - | - | - | - |
| Model I | Lys335 A-Val257 B | Cε-Hε2...O | 2.25 | 0.0130 | 0.0461 | -0.0141 | 17.35 |
| Model II | - | - | - | - | - | - | - |
| Model I | Lys335 A-Val257 B | Nζ-Hζ2...O | 2.38 | 0.0088 | 0.0390 | -0.0141 | 16.58 |
| Model II | - | - | - | - | - | - | - |
| Model I | Ala366 A-Asp256 B | Cβ-Hβ2...O | 2.70 | 0.0053 | 0.0201 | -0.0024 | 4.33 |
| Model II | Ala366 C-Asp256 D |  | 2.57 | 0.0073 | 0.0251 | -0.0041 | 6.31 |
| Model I | Val257 B-Ile364 A | Cγ1-Hγ11...O | 2.69 | 0.0058 | 0.0201 | -0.0027 | 4.60 |
| Model II | Val257 C-Ile364 D |  | 2.67 | 0.0060 | 0.0209 | -0.0029 | 4.87 |
| Model I | Ala293 B-Val330 A | Cα-Hα...O | 2.45 | 0.0104 | 0.0381 | -0.0066 | 9.98 |
| Model II | Ala293 D-Val330 C |  | 2.42 | 0.0102 | 0.0368 | -0.0066 | 9.78 |
| Model I | - | - | - | - | - | - | - |
| Model II | Phe294 D-Val330 C | N-H...O | 1.92 | 0.0243 | 0.0831 | -0.0197 | 26.37 |
| Model I | Glu296 B-Thr329 A | N-H...Oγ1 | 1.81 | 0.0349 | 0.1274 | -0.0295 | 39.75 |
| Model II | Glu296 D-Thr329 C |  | 1.92 | 0.0266 | 0.0844 | -0.0225 | 28.94 |
| Model I | Glu296 B-Thr329 A | Cγ-Hγ2...Oγ1 | 2.60 | 0.0076 | 0.0276 | -0.0043 | 6.76 |
| Model II | - | - | - | - | - | - | - |
| Model I | Gln297 B-Thr328 A | N-H...O | 1.80 | 0.0325 | 0.1146 | -0.0248 | 34.23 |
| Model II | Gln297 D-Thr328 C |  | 1.82 | 0.0310 | 0.1096 | -0.0241 | 33.09 |
| Model I | - | - | - | - | - | - | - |
| Model II | Asn298 D-Thr326 C | Cβ-Hβ3...O | 2.47 | 0.0099 | 0.0361 | -0.0060 | 9.21 |
| Model I | Thr328 B-Gln297 A | N-H...O | 1.80 | 0.0415 | 0.1550 | -0.0345 | 47.15 |
| Model II | Thr328 D-Gln297 C |  | 1.78 | 0.0345 | 0.1278 | -0.0277 | 38.26 |
| Model I | Thr328 B-Gln297 A | Oγ1-Hγ1...O | 2.32 | 0.0101 | 0.0382 | -0.0076 | 10.82 |
| Model II | Thr328 D-Gln297 C |  | 2.41 | 0.0083 | 0.0329 | -0.0056 | 8.53 |
| Model I | Thr329 B-Phe294 A | Cα-Hα...O | 2.60 | 0.0378 | 0.0302 | -0.0100 | 12.08 |
| Model II | Thr329 D-Phe294 C |  | 2.59 | 0.0081 | 0.0282 | -0.0045 | 7.03 |
| Model I | Thr329 B-Phe294 A | Cγ2-Hγ22...O | 2.57 | 0.0091 | 0.0310 | -0.0099 | 12.05 |
| Model II | Thr329 D-Phe294 C |  | 2.63 | 0.0071 | 0.0250 | -0.0038 | 6.04 |
| Model I | Val330 B-Phe294 A | N-H...O | 1.83 | 0.0304 | 0.1085 | -0.0240 | 32.86 |
| Model II | Val330 D-Phe294 C |  | 1.89 | 0.0258 | 0.0894 | -0.0204 | 27.66 |
| Model I | - | - | - | - | - | - | - |
| Model II | Ser331 C-Phe294 D | Cβ-Hβ3...O | 2.34 | 0.0123 | 0.0375 | -0.0090 | 12.02 |
| Model I | Lys335 B-Val257 A | Nζ-Hζ2...O | 2.27 | 0.0136 | 0.0406 | -0.0103 | 13.45 |
| Model II | - | - | - | - | - | - | - |

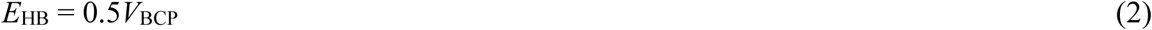

The moduli of the H-bond energies, |*E*_HB_|, of the H-bonds formed in models I and II are also presented in Table 1.

There are two H-bonds of type N-H…O between Gln297 of each subunit and Thr328 of its opposite subunit (Figs 4a and 4b). As is evident in Table 1, all eight H-bonds are moderate H-bonding interactions because their |*E*_HB_| values are located in the range (16.73-62.76 kJ/mol) proposed for moderate H-bonds [9,22]. Among these eight H-bonds, N-H…O in the Thr328 A-Gln297 B pair, wherein Thr328 A acts as a proton donor, is the strongest H-bond because it has the largest values of *ρ*_BCP_ (0.0429 a.u.), ∇^2^*ρ*_BCP_ (0.1648 a.u.), and |*E*_HB_| (49.52 kJ/mol) as well as the smallest length (1.72 Å) among all eight H-bonds. Except the Thr328 C-Gln297 D pair, there is an Oγ1-Hγ1…O H-bond in each Thr328-Gln297 pair. Moreover, Thr328 in both subunits B and D is involved in the Cδ2-Hδ2…O H-bond with Phe294 of subunits A and C, respectively (Fig 4c). Small values of *ρ*_BCP_, ∇^2^*ρ*_BCP_ and |*E*_HB_| in Hγ1…O and Hδ2…O BCPs signify that Oγ1-Hγ1…O and Cδ2-Hδ2…O H-bonds in these residue pairs are weak interactions (Table 1).

**Fig 4.**
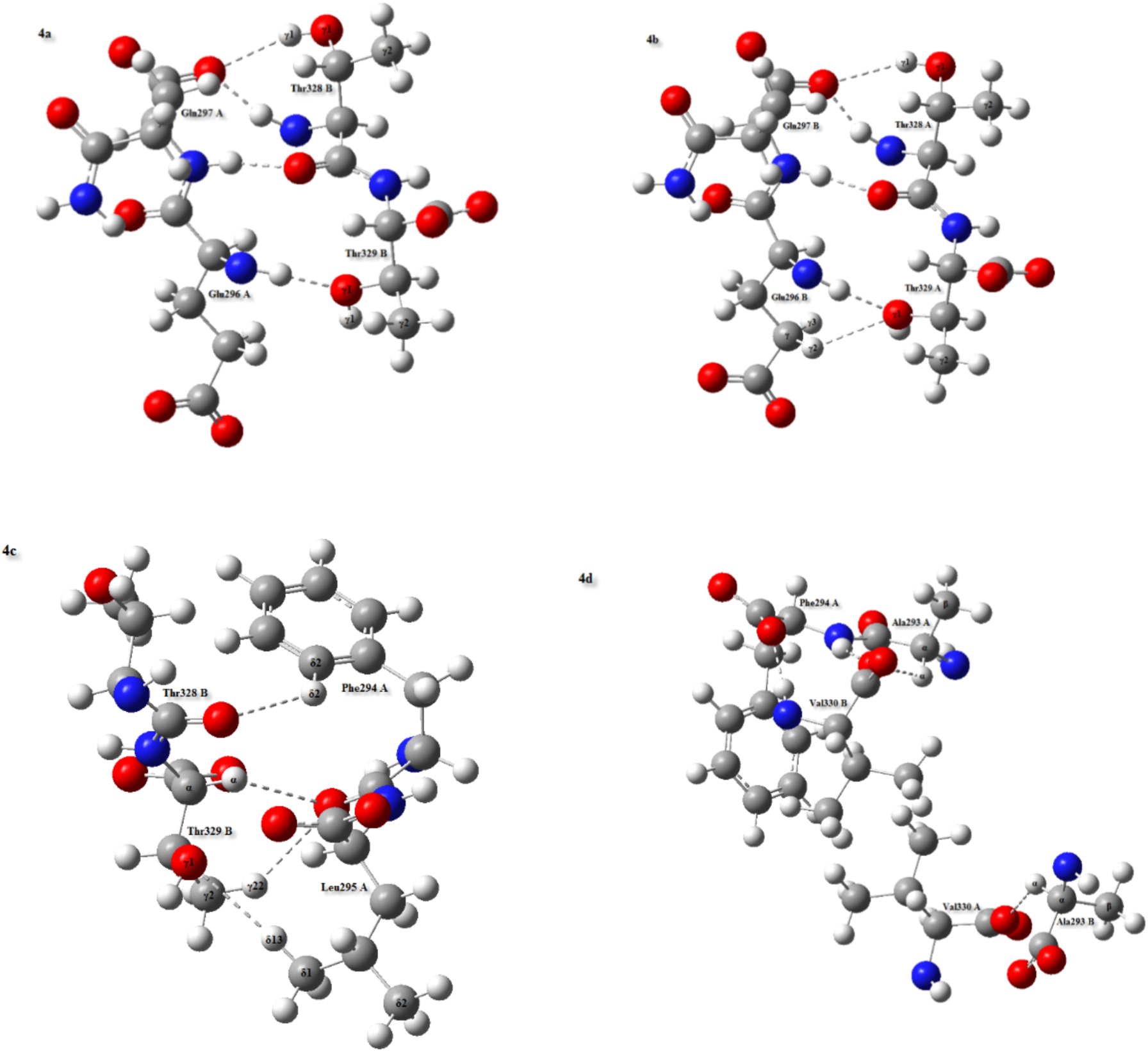
Residue pairs participate in H-bonding interactions at the dimeric A/B interface of the PvNV tetrameric P-domain. These H-bonds are repeated between each corresponding residue pair in its dimeric C/D interface. The oxygen, carbon, nitrogen and hydrogen atoms are shown in red, grey, blue and white colors, respectively.

QTAIM analysis detects that Phe294 of subunits A and C can also form Hα…O and Hγ22…O BCPs with Thr329 of subunits B and D, respectively. Since Cα-Hα…O and Cγ2-Hγ22…O H-bonds in the Thr329 B-Phe294 A pair have larger values of |*E*_HB_| than for the Thr329 D-Phe294 C pair, these H-bond strengths in the former pair are greater than in the latter pair. As shown in Table 1, with the exception of the Phe294 B-Val330 A pair, Phe294 of each subunit also participates in two H-bonding interactions of type N-H…O with Val330 from its contrary subunit (Fig 4d). Among these six moderate H-bonds, the largest values of *ρ*_BCP_ (0.0302 a.u.), ∇^2^*ρ*_BCP_ (0.1137 a.u.) and |*E*_HB_| (33.41 kJ/mol) belong to the N-H…O H-bond in the Phe294 A-Val330 B pair in which Phe294 A behaves as a proton donor. This H-bond is thus the strongest of the six H-bonds. Additionally, a weak H-bond of Cβ-Hβ3…O with |*E*_HB_| 12.02 kJ/mol and length 2.34 Å is found between Phe294 D and Ser331 C.

QTAIM analysis identifies the presence of a Hα…O BCP in each Ala293-Val330 pair, except the Ala293 C-Val330 D pair. The values of |*E*_HB_| for Cα-Hα…O in Ala293 B-Val330 A and Ala293 D-Val330 C pairs are 9.98 and 9.78 kJ/mol, respectively; their strengths in these two pairs are thus the same. As presented in Table 1, there is one BCP that connects the Oγ1 nucleus of Thr329 of each subunit to the hydrogen nucleus of Glu296 from its opposite subunit. The topological features of H…Oγ1 BCPs confirm that Glu296-Thr329 pairs are able to form stronger N-H…Oγ1 H-bonds in model I than in model II. In addition to this H-bond, Glu296 B interacts with Thr329 A through a weak H-bond of Cγ-Hγ2…Oγ1 (Fig 4b). Furthermore, the Hδ13…Oγ1 BCPs links the Oγ1 nuclei in Thr329 B and Thr329 D to the Hδ13 nuclei in Leu295 A and Leu295 C, respectively. Two H-bonds of Cδ1-Hδ13…Oγ1 in Leu295 A-Thr329 B and Leu295 C-Thr329 D pairs are weak interactions, reflecting the values of their topological parameters (Table 1).

QTAIM analysis indicates that the O nucleus of Val257 B is simultaneously connected to Hε2 and Hζ2 of Lys335 A with two BCP. As |*E*_HB_| of Cε-Hε2…O (17.35 kJ/mol) is only slightly greater than that of Nζ-Hζ2…O (16.58 kJ/mol), the unconventional H-bond is slightly stronger than the conventional H-bond. Although Nζ-Hζ2…O is formed between Val257 A and Lys335 B, its strength becomes less in this pair through a smaller |*E*_HB_| (13.45 kJ/mol). Furthermore, there is a BCP between Hζ1 of Lys335 A and Oδ1 of Asp256 B. Large values of *ρ*_BCP_ (0.0588) and ∇^2^*ρ*_BCP_ (0.1466) of Hζ1…Oδ1 BCP demonstrate that Nζ-Hζ1…Oδ1 in the Lys335 A-Asp256 B pair is a strong H-bond. This H-bond with |*E*_HB_| 64.81 kJ/mol and length 1.61 Å is the strongest H-bonding interaction within the dimeric interfaces of the tetrameric conformation of PvNVPd. It is reasonable to suggest that the Nζ-Hζ1…Oδ1 H-bond plays an essential role in the stability of the dimeric interface between subunits A and B. It is worth mentioning that no H-bond formed in three pairs of Lys335 A-Asp256 B, Lys335 A-Val257 B and Lys335 B-Val257 A is characterized in similar residue pairs inside the dimeric interface between subunits C and D (Fig 5 and Table 1).

**Fig 5.**
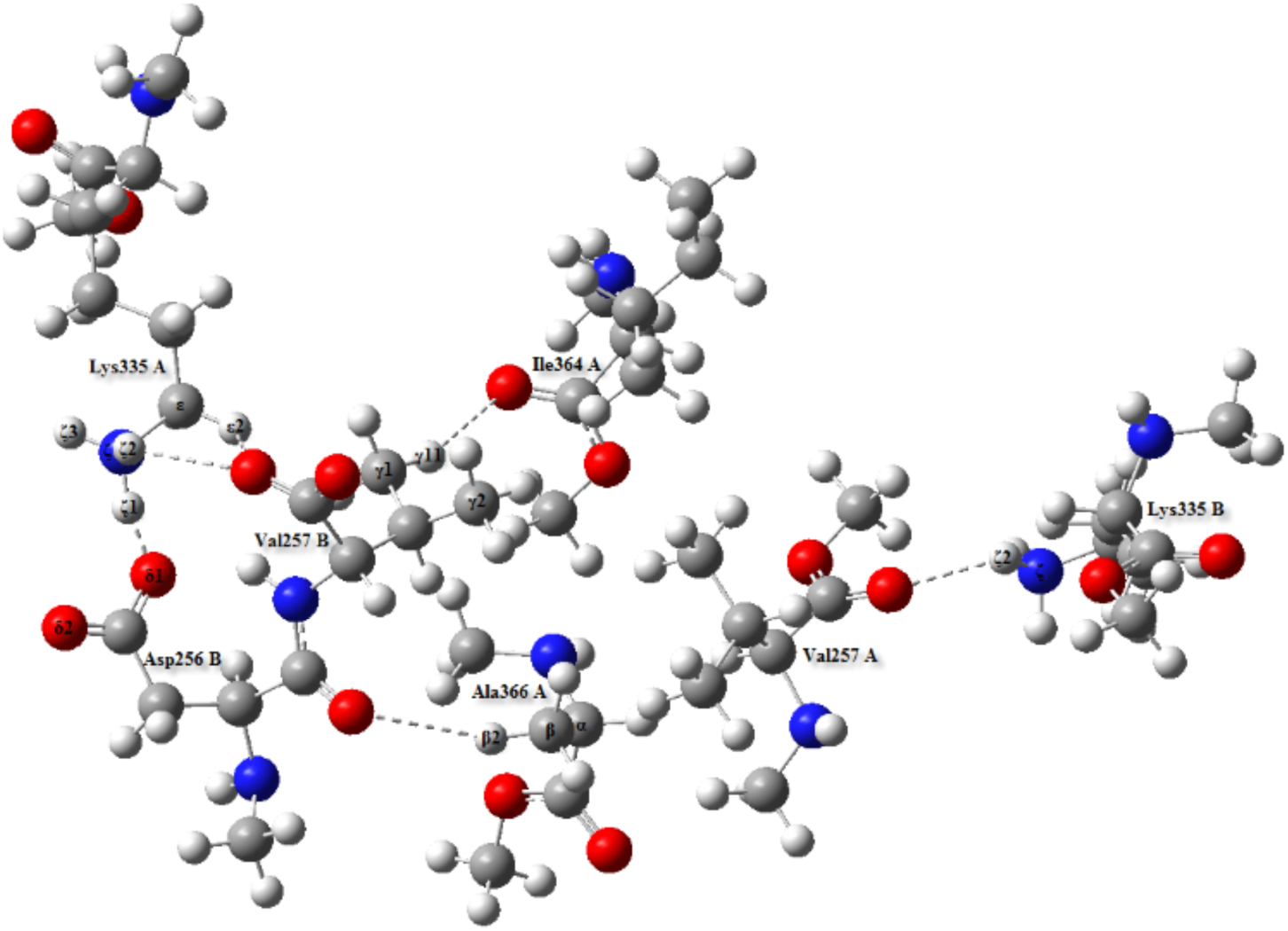
H-bonds formed in Lys335 A-Asp256 B, Lys335 A-Val257 B, Lys335 B-Val257 A, Ala366 A-Asp256 B and Val257 B-Ile364 A pairs.

As seen in Table 1, there are two weak H-bonds of Cγ1-Hγ11…O in Val257 B-Ile364 A and Val257 C-Ile364 D pairs as well as two weak H-bond of Cβ-Hβ2…O detected in Ala366 A-Asp256 B and Ala366 C-Asp256 D pairs (Fig 5). Fig 6 displays that Thr326 C is involved in two weak H-bonds of Cα-Hα…Oδ1 and Cβ-Hβ3…O with Asn298 D. Likewise, Pro325 C interacts with Gln300 D through a weak H-bond of Cβ-Hβ2…Oε1. These three H-bonds are not detected in two pairs of Thr326 A-Asn298 B and Pro325 A-Gln300 B.

**Fig 6.**
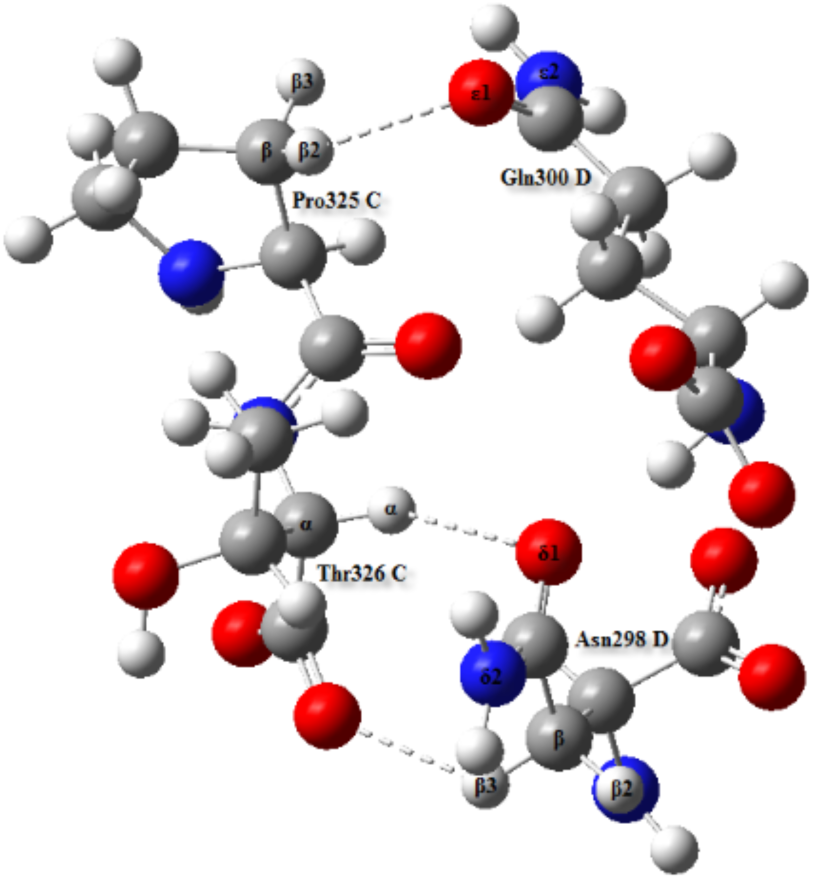
Residues Pro325 and Thr326 of subunit C interact with residues Gln300 and Asn298 of subunit D, respectively.

According to the Rozas criteria [27], positive ∇^2^*ρ*_BCP_ and negative *H*_BCP_ values in H…B BCP are characteristic of a partially covalent nature in a pertinent H-bonding interaction. It is evident from the results in Table 1 that all discussed H-bonding interactions have positive values of ∇^2^*ρ*_BCP_ and negative values of *H*_BCP_. The fundamental nature of these interactions must therefore be considered as intermediate between covalent and electrostatic character. Based on the *ρ*_BCP_ values, a strong H-bond of Nζ-Hζ1…Oδ1 in Lys335 A-Asp256 B pair has a basically covalent nature, whereas all weak H-bonds in both models have mainly an electrostatic character. Moderate H-bonds are a mixture of both covalent and electrostatic contributions in that their covalent nature is decreased on decreasing their strengths. Based on the magnitude of |*E*_HB_|, the covalent part reduction of the putative H-bonds in the interacting residue pairs of both models has the following order: Nζ-Hζ1…Oδ1 (Lys335 A-Asp256 B) > N-H…O (Thr328 A-Gln297 B) > N-H…O (Thr328 B-Gln297 A) > N-H…Oγ1 (Glu296 B-Thr329 A) > N-H…Oγ1 (Glu296 A-Thr329 B) > N-H…O (Thr328 D-Gln297 C) > N-H…O (Gln297 A-Thr328 B) ≅ N-H…O (Gln297 B-Thr328 A) > N-H…O (Phe294 A-Val330 B) ≅ N-H…O (Gln297 D-Thr328 C) > N-H…O (Val330 B-Phe294 A) > N-H…O (Thr328 C-Gln297 D) > N-H…Oγ1 (Glu296 C-Thr329 D) > N-H…O (Gln297 C-Thr328 D) > N-H…Oγ1 (Glu296 D-Thr329 C) > N-H…O (Val330 D-Phe294 C) > N-H…O (Phe294 D-Val330 C) > N-H…O (Val330 C-Phe294 D) > N-H…O (Phe294 C-Val330 D) > Cε-Hε2…O (Lys335 A-Val257 B) > Nζ-Hζ2…O (Lys335 A-Val257 B) > Nζ-Hζ2…O (Lys335 B-Val257 A) > Cα-Hα…O (Thr329 B-Phe294 A) ≅ Cγ2-Hγ22…O (Thr329 B-Phe294 A) ≅ Cβ-Hβ3…O (Ser331 C-Phe294 D) ≅ Cα-Hα…O (Ala293 A-Val330 B) > Oγ1-Hγ1…O (Thr328 B-Gln297 A) > Cα-Hα…O (Ala293 B-Val330 A) ≅ Cα-Hα…O (Ala293 D-Val330 C) > Cβ-Hβ3…O (Asn298 D-Thr326 C) > Oγ1-Hγ1…O (Thr328 A-Gln297 B) ≅ Cδ1-Hδ13…Oγ1 (Leu295 C-Thr329 D) ≅ Oγ1-Hγ1…O (Thr328 D-Gln297 C) > Cδ1-Hδ13…Oγ1 (Leu295 A-Thr329 B) ≅ Cα-Hα…O (Thr329 D-Phe294 C) > Cα-Hα…Oδ1 (Thr326 C-Asn298 D) ≅ Cγ-Hγ2…Oγ1 (Glu296 B-Thr329 A) > Cδ2-Hδ2…O (Phe294 A-Thr328 B) ≅ Cβ-Hβ2…O (Ala366 C-Asp256 D) > Cγ2-Hγ22…O (Thr329 D-Phe294 C) ≅ Cδ2-Hδ2…O (Phe294 C-Thr328 D) > Cβ-Hβ2…Oε1 (Pro325 C-Gln300 D) > Cγ1-Hγ11…O (Val257 C-Ile364 D) ≅ Cγ1-Hγ11…O (Val257 B-Ile364 A) ≅ Cβ-Hβ2…O (Ala366 A-Asp256 B).

The above trend reveals that the length of the H-bond and the |*E*_HB_| are two effective factors pertaining to the strength of a given H-bond. Our results indicate the existence of a linear correlation between the H-bond distances and the estimated ln |*E*_HB_| in both models I and II described with the following regression equation (Fig 7):

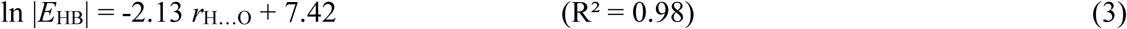

**Fig 7.**
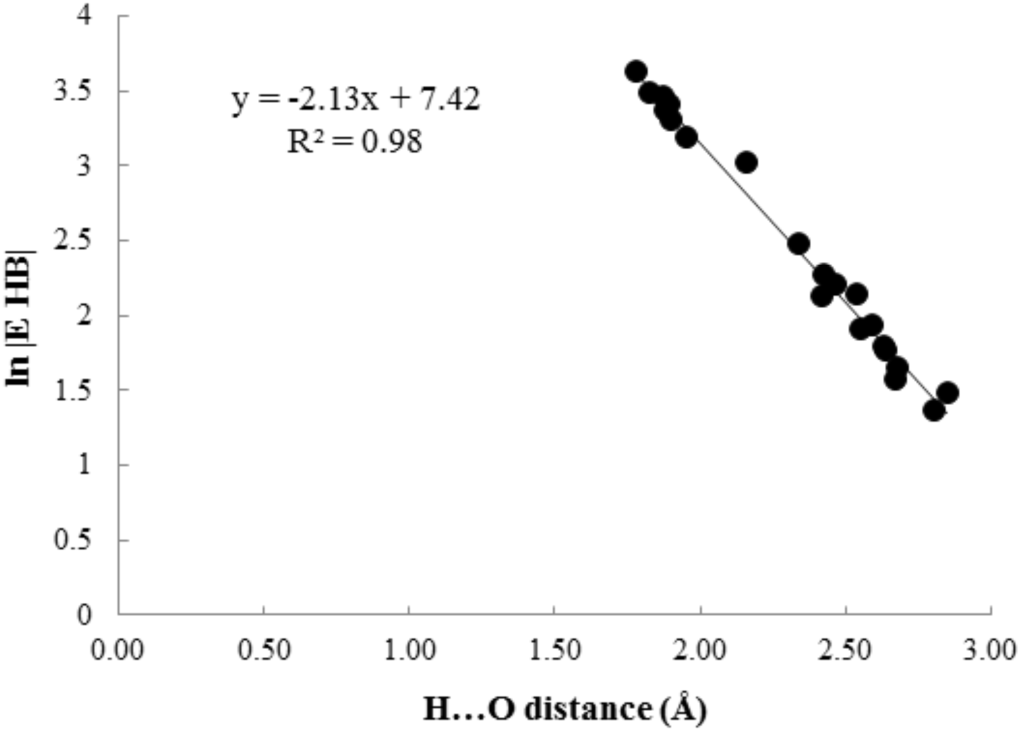
Correlation between ln |*E*_HB_| and putative H-bond length in the dimeric C/D interface (model II). The negative slope means that there is an inverse relation between the strength and the length of a pertinent H-bond; a shorter H-bond is hence accompanied with a stronger H-bonding interaction.

### QTAIM analysis on the parallel dimeric interface of PvNVPd

The QTAIM analysis of the parallel dimeric A/B interface demonstrates that the type of H-bond formed in model III are common to those in models I and II (Table 2).

**Table 2.** H-bond distance (*d*) and selected topological parameters of electron density in various H…B BCP characterized inside the dimeric A/B interface of the parallel conformation (model III), evaluated at the M06-2X/6-31G** level.

| Residue pair | H-bond | $d$<br>(Å) | $\rho_{\text{BCP}}$<br>(a.u.) | $\nabla^2\rho_{\text{BCP}}$<br>(a.u.) | $H_{\text{BCP}}$<br>(a.u.) | $\vert E_{\text{HB}}\vert $<br>(kJ/mol) |
| --- | --- | --- | --- | --- | --- | --- |
| Ala293 A-Val330 B | Cα-Hα...O | 2.43 | 0.0105 | 0.0382 | -0.0066 | 9.99 |
| Phe294 A-Thr328 B | Cδ2-Hδ2...O | 2.72 | 0.0058 | 0.0221 | -0.0027 | 4.76 |
| Phe294 A-Thr328 B | Cε-Hε2...O | 2.69 | 0.0069 | 0.0243 | -0.0034 | 5.62 |
| Phe294 A-Val330 B | N-H...O | 1.93 | 0.0235 | 0.0833 | -0.0191 | 25.79 |
| Leu295 A-Thr329 B | Cδ1-Hδ13...Oγ1 | 2.77 | 0.0059 | 0.0200 | -0.0027 | 4.59 |
| Glu296 A-Thr329 B | N-H...Oγ1 | 1.91 | 0.0269 | 0.0863 | -0.0223 | 28.99 |
| Gln297 A-Thr328 B | N-H...O | 1.90 | 0.0261 | 0.0870 | -0.0209 | 27.78 |
| Asn298 A-Thr326 B | Cβ-Hβ3...O | 2.37 | 0.0119 | 0.0410 | -0.0080 | 11.49 |
| Thr328 A-Gln297 B | N-H...O | 1.73 | 0.0381 | 0.1396 | -0.0298 | 41.35 |
| Thr328 A-Gln297 B | Oγ1-Hγ1...O | 2.53 | 0.0065 | 0.0279 | -0.0039 | 6.50 |
| Thr329 A-Phe294 B | Cγ2-Hγ22...O | 2.61 | 0.0076 | 0.0261 | -0.0042 | 6.52 |
| Val330 A-Phe294 B | N-H...O | 2.00 | 0.0210 | 0.0684 | -0.0174 | 22.74 |
| Ser331 A-Phe294 B | Cβ-Hβ3...O | 2.34 | 0.0120 | 0.0367 | -0.0088 | 11.69 |
| Lys335 A-Asp256 B | Nζ-Hζ1...Oδ1 | 1.61 | 0.0588 | 0.1467 | -0.0557 | 64.76 |
| Lys335 A-Val257 B | Cε-Hε2...O | 2.44 | 0.0088 | 0.0390 | -0.0054 | 9.00 |
| Lys335 A-Val257 B | Nζ-Hζ2...O | 2.38 | 0.0130 | 0.0461 | -0.0090 | 12.92 |
| Ala293 B-Val330 A | Cα-Hα...O | 2.49 | 0.0090 | 0.0333 | -0.0054 | 8.38 |
| Phe294 B-Thr328 A | Cε-Hε2...O | 2.69 | 0.0071 | 0.0247 | -0.0035 | 5.76 |
| Phe294 B-Val330 A | N-H...O | 1.94 | 0.0232 | 0.0792 | -0.0190 | 25.30 |
| Leu295 B-Thr329 A | Cδ1-Hδ13...Oγ1 | 2.71 | 0.0063 | 0.0226 | -0.0033 | 5.33 |
| Glu296 B-Thr329 A | N-H...Oγ1 | 1.89 | 0.0290 | 0.0915 | -0.0240 | 31.00 |
| Glu296 B-Thr329 A | Cγ-Hγ2...Oγ1 | 2.70 | 0.0066 | 0.0238 | -0.0035 | 5.68 |
| Gln297 B-Thr328 A | N-H...O | 1.90 | 0.0265 | 0.0887 | -0.0212 | 28.28 |
| Thr328 B-Gln297 A | N-H...O | 1.85 | 0.0303 | 0.1004 | -0.0239 | 31.87 |
| Thr328 B-Gln297 A | Oγ1-Hγ1...O | 2.52 | 0.0064 | 0.0275 | -0.0039 | 6.40 |
| Thr329 B-Phe294 A | Cγ2-Hγ22...O | 2.60 | 0.0076 | 0.0265 | -0.0043 | 6.62 |
| Val330 B-Phe294 A | N-H...O | 1.99 | 0.0216 | 0.0697 | -0.0178 | 23.19 |

In model III, conformational changes lead to decrease values of *ρ*_BCP_, ∇^2^*ρ*_BCP_ and |*E*_HB_| of Hα…O, Hδ2…O, H…O, Hγ1…O, H…O, and Hδ13…Oγ1 BCPs in Ala293-Val330, Phe294 A-Thr328 B, Phe294-Val330, Thr328-Gln297, Gln297-Thr328, and Leu295-Thr329 pairs, respectively (Tables 1 and 2). H-bonds Cα-Hα…O, Cδ2-Hδ2…O, N-H…O, Oγ1-Hγ1…O, N-H…O and Cδ1-Hδ13…Oγ1 in the cited pairs are therefore converted to interactions weaker than those formed in models I and II. Among them, when Thr328 A acts as a proton donor, its N-H…O H-bond with Gln297 B is the strongest because values of *ρ*_BCP_ (0.0381 a.u.), ∇^2^*ρ*_BCP_ (0.1396 a.u.) and |*E*_HB_| (41.35 kJ/mol) of this H-bond are the largest. The values of |*E*_HB_| of Cγ2-Hγ22…O H-bonds in Thr329 A-Phe294 B, Thr329 B-Phe294 A and Thr329 D-Phe294 C pairs are 6.52, 6.62 and 6.04 kJ/mol, respectively; these three weak interactions have hence equal strengths. Likewise, a weak Cβ-Hβ3…O H-bond in the Ser331 A-Phe294 B pair is topologically equivalent to this H-bond in the Ser331 C-Phe294 D pair (Tables 1 and 2).

Similar to model I, Nζ-Hζ1…Oδ1 in the Lys335 A-Asp256 B pair has a major contribution to the dimeric interactions of the dimeric interface of parallel PvNVPd because this H-bond with |*E*_HB_| 64.76 and length 1.61 Å is the strongest H-bonding interaction in model III (Table 2). Conformational changes convert two H-bonds of Cε-Hε2…O and Nζ-Hζ2…O in the Lys335 A-Val257 B pair from moderate interactions in model I to weak interactions in model III. In contrast, the values of *ρ*_BCP_, ∇^2^*ρ*_BCP_, and |*E*_HB_| of Hβ3…O BCP in Asn298 A-Thr326 B and Asn298 D-Thr326 C pairs indicate that Cβ-Hβ3…O is stronger H-bond in model III than in model II. The strengths of the N-H…Oγ1 H-bonds in Glu296 B-Thr329 A and Glu296 C-Thr329 D pairs are nearly identical because |*E*_HB_| (31.00 kJ/mol) in model III is almost equal to that (30.43 kJ/mol) in model II.

Beyond the mentioned H-bonds, there are two weak H-bonds of Cε-Hε2…O in Phe294 A-Thr328 B and Phe294 B-Thr328 A pairs, neither of which is observed in Phe294-Thr328 pairs of models I and II. Similarly, we found a linear correlation between the lengths of H-bonds and the estimated ln |*E*_HB_| in model III (ln |*E*_HB_| = −2.15 *r*_H…O_ + 7.47) with linear correlation coefficient 0.993. The magnitude |*E*_HB_| predicts that the strength of the putative H-bonds in the interacting residue pairs of model III decreases in the following order: Nζ-Hζ1…Oδ1 (Lys335 A-Asp256 B) > N-H…O (Thr328 A-Gln297 B) > N-H…O (Thr328 B-Gln297 A) > N-H…Oγ1 (Glu296 B-Thr329 A) > N-H…Oγ1 (Glu296 A-Thr329 B) > N-H…O (Gln297 B-Thr328 A) ≅ N-H…O (Gln297 A-Thr328 B) > N-H…O (Phe294 A-Val330 B) ≅ N-H…O (Phe294 B-Val330 A) > N-H…O (Val330 B-Phe294 A) ≅ N-H…O (Val330 A-Phe294 B) >Nζ-Hζ2…O (Lys335 A-Val257 B) > Cβ-Hβ3…O (Ser331 A-Phe294 B) ≅ Cβ-Hβ3…O (Asn298 A-Thr326 B) > Cα-Hα…O (Ala293 A-Val330 B) > Cε-Hε2…O (Lys335 A-Val257 B) > Cα-Hα…O (Ala293 B-Val330 A) > Cγ2-Hγ22…O (Thr329 B-Phe294 A) ≅ Cγ2-Hγ22…O (Thr329 A-Phe294 B) ≅ Oγ1-Hγ1…O (Thr328 A-Gln297 B) ≅ Oγ1-Hγ1…O (Thr328 B-Gln297 A) > Cε-Hε2…O (Phe294 B-Thr328 A) ≅ Cγ-Hγ2…Oγ1 (Glu296 B-Thr329 A) ≅ Cε-Hε2…O (Phe294 A-Thr328 B) ≅ Cδ1-Hδ13…Oγ1 (Leu295 B-Thr329 A) ≅ Cδ2-Hδ2…O (Phe294 A-Thr328 B) ≅ Cδ1-Hδ13…Oγ1 (Leu295 A-Thr329 B).

This QTAIM analysis evaluates that the total hydrogen bond energies at the BCP detected in models I, II and III are 513.32, 386.37 and 472.27 kJ/mol, respectively. The total H-bonding interactions present in model I are consequently stronger than those formed in models II and III.

### NBO analysis of the dimeric interfaces of the tetrameric conformation of PvNVPd

The QTAIM analysis identified the conventional and unconventional H-bonding interactions of various types contributing to the dimeric interactions at the dimeric interfaces of PvNVPd in the tetrameric conformation. In this section, the strengths of the local orbitals participating in each of these H-bonds are evaluated with the second-order perturbation energies resulted from the NBO analysis (Table 3).

**Table 3.**
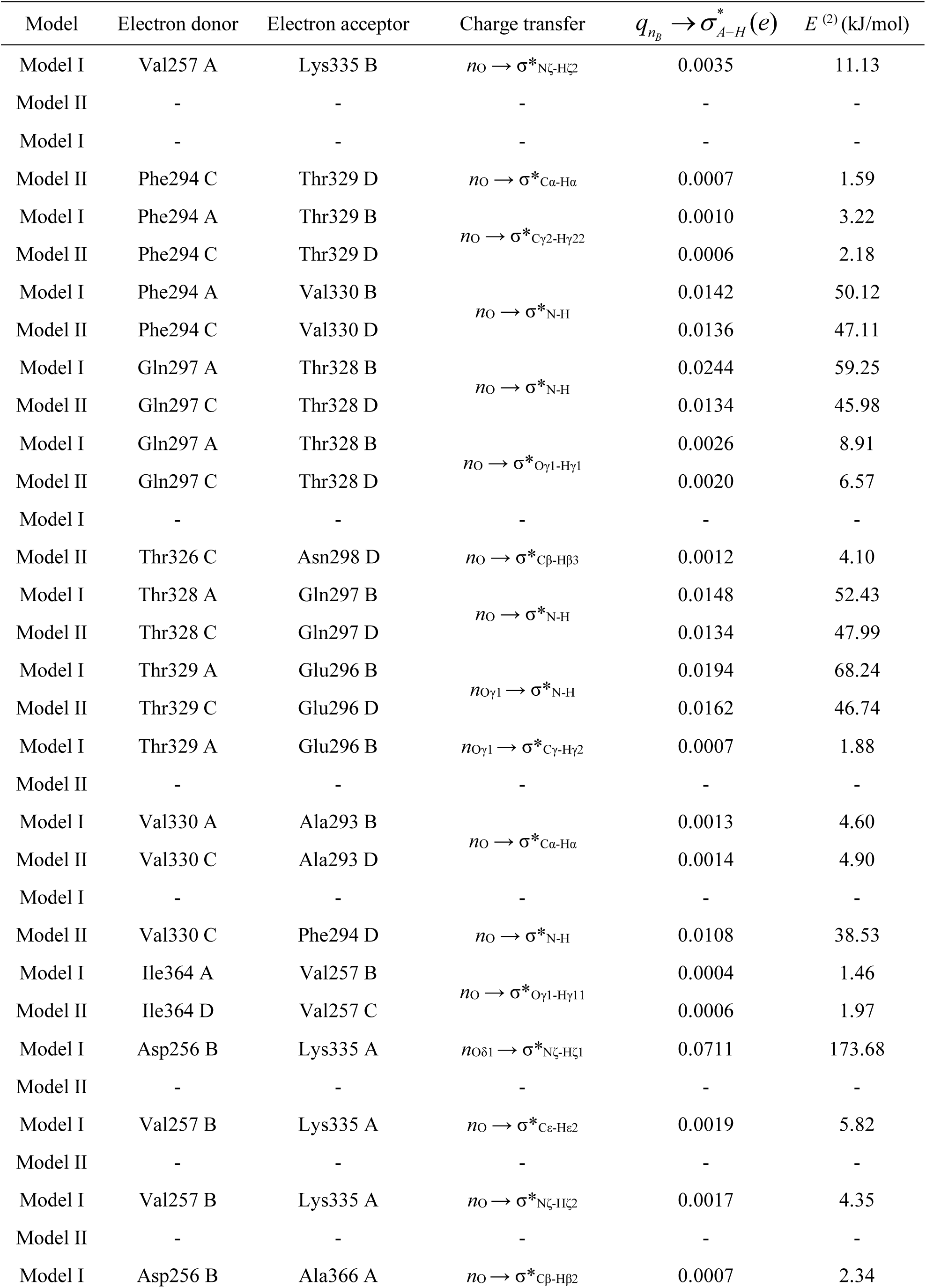
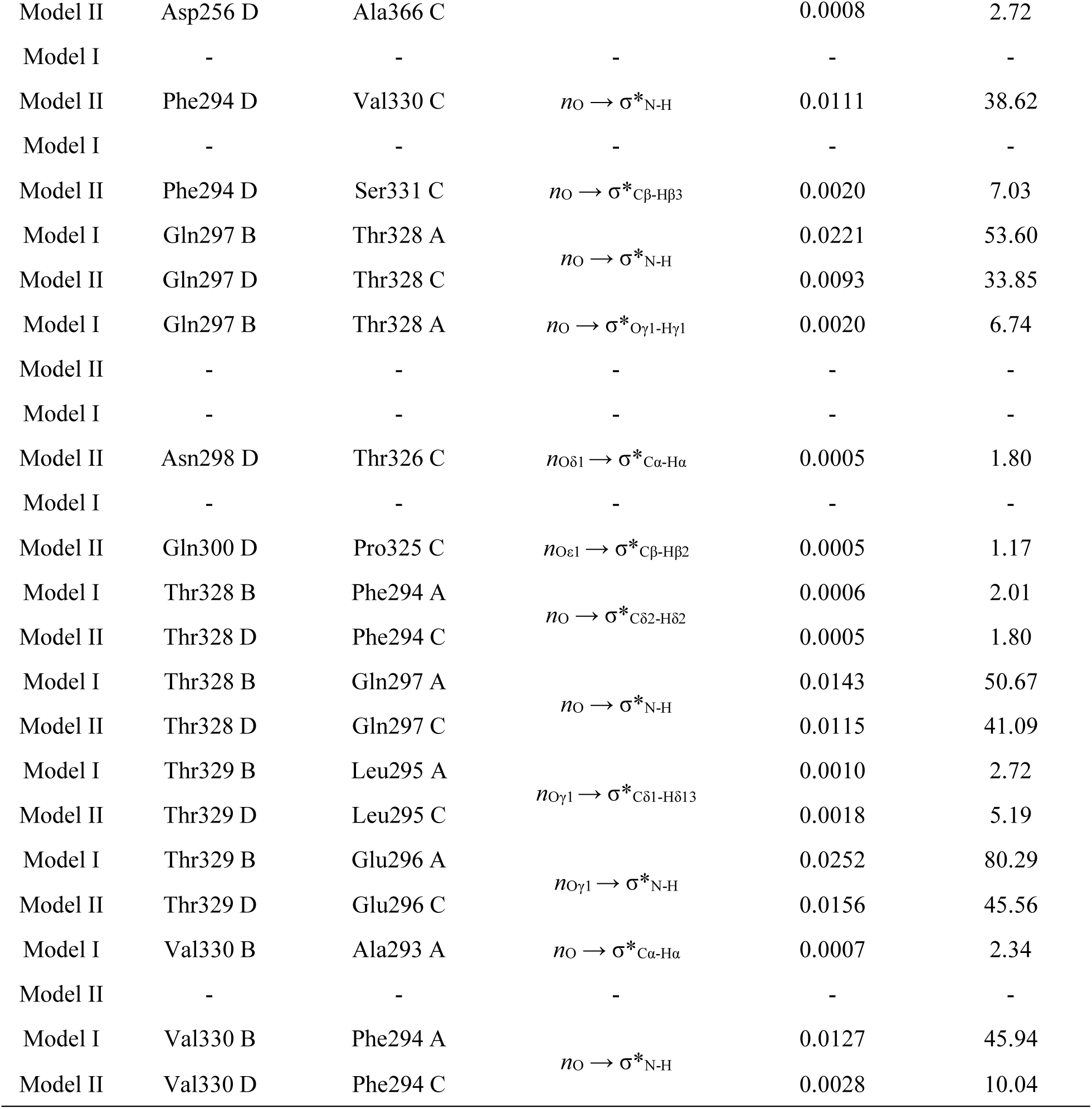
NBO results of local orbitals of partner atoms in charge-transfer interactions within the dimeric A/B (model I) and C/D (model II) interfaces of the tetrameric protein, calculated at the M06-2X/6-31G** level.

The NBO analysis shows the presence of two charge transfer interactions of type *n*_O_ → σ*_N-H_ between Gln297 of each subunit and Thr328 from its inverse subunit. Among them, the highest values of *q*_CT_ (0.0244 *e*) and *E* ^(2)^ (59.25 kJ/mol) pertain to the *n*_O_ → σ*_N-H_ interaction of the Gln297 A-Thr328 B pair wherein Gln297 A behaves as an electron donor (Table 3). Although this interaction is the strongest *n*_O_ → σ*_N-H_ interaction in the Gln297-Thr328 pairs, it is not the strongest H-bonding interaction in these residue pairs. The reason for this discrepancy is that the QTAIM analysis established that N-H…O arising from this charge-transfer interaction with |*E*_HB_| 47.15 kJ/mol and length 1.80 Å is the second strongest H-bond in the Gln297-Thr328 pairs (Table 1). NBO analysis predicts that the strongest N-H…O existing in a Gln297 B-Thr328 A pair is the result of charge transfer (0.0221 e) from *n*_O_ of Gln297 B to σ*_N-H_ of Thr328 A with *E* ^(2)^ of 53.60 kJ/mol. This charge transfer is the second strongest *n*_O_ → σ*_N-H_ interaction in the Gln297-Thr328 pairs.

The *n*_O_ lone pairs of Gln297 of subunits A, B and C also interact weakly with the antibonding orbitals of σ*_Oγ1-Hγ1_ in Thr328 of subunits B, A and D, respectively. Besides, two weak interactions of type *n*_O_ → σ*_Cδ2-Hδ2_ exist in Phe294 A-Thr328 B and Phe294 C-Thr328 D pairs. *n*_O_ of Phe294 A overlaps simultaneously with the antibonding orbitals of σ*_Cγ2-Hγ22_ in Thr329 B and σ*_N-H_ in Val330 B. *E* ^(2)^ of the *n*_O_ → σ*_N-H_ (50.12 kJ/mol) is more than fifteen times that of the *n*_O_ → σ*_Cγ2-Hγ22_ interaction (3.22 kJ/mol). Phe294 A has thus a much stronger donor-acceptor interaction with Val330 B than with Thr329 B. In contrast, there is an orbital overlap between σ*_N-H_ of Phe294 A and *n*_O_ of Val330 B but with a smaller *E* ^(2)^ (45.94 kJ/mol). Similarly, two interactions of type *n*_O_ → σ*_N-H_ occur in each Phe294-Val330 pair of model II; a weak interaction of *n*_O_ → σ*_Cγ2-Hγ22_ is repeated in the Phe294 C-Thr329 D pair. The charge transfer (0.0136 *e*) from *n*O of Phe294 C to σ*_N-H_ of Val330 D with *E* ^(2)^ 47.11 kJ/mol is the strongest *n*O → σ*_N-H_ interaction in Phe294-Val330 pairs of model II (Table 3).

An infinitesimal charge transfer (0.0007 *e*) occurs from *n*_O_ of Val330 B to σ*_Cα-Hα_ in Ala293 A with stabilization energy 2.34 kJ/mol. The *n*_O_ → σ*_Cα-Hα_ interactions appear also in Ala293 B-Val330 A and Ala293 D-Val330 C pairs. The donor-acceptor interactions in these two residue pairs have the same strengths, reflecting their equal *E*^(2)^ values (Table 3). The NBO analysis reveals that *n*_Oγ1_ of Thr329 of each subunit donates its electrons to σ*_N-H_ of Glu296 from its counter subunit. The amounts of *E*^(2)^ and *q*_CT_ of the *n*_Oγ1_ → σ*_N-H_ interactions demonstrate that Glu296-Thr329 pairs of model I have stronger donor-acceptor interactions than those of model II; this finding agrees with the |*E*_HB_| prediction. Of these, the *n*_Oγ1_ → σ*_N-H_ interaction in the Glu296 A-Thr329 B pair with *q*_CT_ 0.0252 *e* and *E*^(2)^ 80.29 kJ/mol is the strongest among the four interactions. The *n*_Oγ1_ lone pairs of Thr329 B and Thr329 D overlap weakly with the antibonding orbitals of σ*_Cδ1-Hδ13_ in Leu295 A and Leu295 C, respectively, resulting in the formation of weak H-bonds of Cδ1-Hδ13…Oγ1 in these pairs. Besides, weak Cγ-Hγ2…Oγ1 in Glu296 B-Thr329 A pair is the consequence of a very weak attractive interaction between *n*_Oγ1_ of Thr329 A and the σ*_Cγ-Hγ2_ of Glu296 B with *E*^(2)^ 1.88 kJ/mol.

NBO analysis shows the appearance of two interactions of *n*_O_ → σ*_Cβ-Hβ3_ and *n*_Oδ1_ → σ*_Cα-Hα_ in the Thr326 C-Asn298 D pair as well as *n*_O_ → σ*_Cβ-Hβ3_ and *n*_Oε1_ → σ*_Cβ-Hβ2_ interactions in the Ser331 C-Phe294 D and Pro325 C-Gln300 D pairs, respectively. All five charge transfers are weak interactions, reflecting very small values of their *q*_CT_ and *E*^(2)^ (Table 3). None of these interactions is observed in the corresponding pairs of model I. Based on the |*E*_HB_| results, Nζ-Hζ1…Oδ1 in the Lys335 A-Asp256 B pair is the strongest H-bond in both models. This H-bond is the result of a great charge transfer (0.0711 *e*) from *n*_Oδ1_ of Asp256 B to σ*_Nζ-Hζ1_ of Lys335 A. The *n*_Oδ1_ → σ*_Nζ-Hζ1_ interaction with *E*^(2)^ 173.68 kJ/mol is the strongest charge-transfer interaction in both models. Thereby the NBO results also confirm the great importance of this interaction in maintaining the dimeric interface between subunits A and B. Additionally, there are weak interactions of *n*_O_ → σ*_Cε-Hε2_ and *n*_O_ → σ*_Nζ-Hζ2_ in the Lys335 A-Val257 B and Lys335 B-Val257 A pairs. These interactions do not occur between the corresponding residues in model II (Table 3).

Cε-Hε2…O in the Lys335 A-Val257 B pair is the consequence of a charge transfer (0.0019 *e*) from *n*_O_ of Val257 B to σ*_Cε-Hε2_ of Lys335 A. As seen in Table 1, it is the strongest H-bond of type C-H…O in both models because its topological parameters as well as its stabilization energy (5.82 kJ/mol) are the largest among unconventional H-bonds. Our NBO results show that the *n*_O_ → σ*_Oγ1-Hγ11_ interaction in the Val257 B-Ile364 A pair is the weakest charge-transfer interaction in model I because it has the smallest stabilization energy (1.46 kJ/mol) in comparison with the other interactions.

### NBO analysis on the parallel dimeric interface of PvNVPd

QTAIM analysis established that the N-H…O H-bonds in the Phe294-Val330 and Gln297-Thr328 pairs become weaker in model III. The reason is that each *n*_O_ → σ*_N-H_ interactions in each residue pairs emerged with smaller values of *q*_CT_ and *E*^(2)^ in model III than in models I and II (Table 4). Two interactions of *n*_O_ → σ*_Cγ2-Hγ22_ in Phe294-Thr329 pairs of model III have strengths equivalent with this interaction in the Phe294 A-Thr329 B pair of model I, reflecting the same values of their *q*_CT_ and *E*^(2)^. The stabilization energy of each *n*_O_ → σ*_Oγ1-Hγ1_ interaction in each Gln297-Thr328 pair of model I is about twice that in the corresponding pair of model III; these interaction strengths in model I are hence more than in model III. Similar to models I and II, two interactions of *n*_O_ → σ*_Cε-Hε2_ and *n*_O_ → σ*_Nζ-Hζ2_ in the Lys335 A-Val257 B pair as well as the interactions of *n*_O_ → σ*_Cα-Hα_, *n*_Oγ1_ → σ*_Cδ1-Hδ13_, *n*_O_ → σ*_Cβ-Hβ3_, and *n*_Oγ1_ → σ*_Cγ-Hγ2_ in Ala293-Val330, Thr329-Leu295, Ser331 A-Phe294 B and Glu296 B-Thr329 A pairs, respectively, are weak donor-acceptor interactions (Tables 3 and 4).

**Table 4.** NBO results of local orbitals of partner atoms in charge-transfer interactions inside the dimeric A/B interface of the parallel conformation (model III), estimated at the M06-2X/6-31G** level.

| Electron donor | Electron acceptor | Charge transfer | $q_{n_B} \rightarrow \sigma^*_{A-H}(e)$ | $E^{(2)}$ (kJ/mol) |
| --- | --- | --- | --- | --- |
| Phe294 A | Thr329 B | $n_{\text{O}} \rightarrow \sigma^*_{\text{C}\gamma 2\text{-H}\gamma 22}$ | 0.0010 | 3.18 |
| Phe294 A | Val330 B | $n_{\text{O}} \rightarrow \sigma^*_{\text{N-H}}$ | 0.0088 | 30.71 |
| Gln297 A | Thr328 B | $n_{\text{O}} \rightarrow \sigma^*_{\text{N-H}}$ | 0.0119 | 41.25 |
| Gln297 A | Thr328 B | $n_{\text{O}} \rightarrow \sigma^*_{\text{O}\gamma 1\text{-H}\gamma 1}$ | 0.0013 | 4.27 |
| Thr328 A | Gln297 B | $n_{\text{O}} \rightarrow \sigma^*_{\text{N-H}}$ | 0.0115 | 40.84 |
| Thr329 A | Leu295 B | $n_{\text{O}\gamma 1} \rightarrow \sigma^*_{\text{C}\delta 1\text{-H}\delta 13}$ | 0.0011 | 3.26 |
| Thr329 A | Glu296 B | $n_{\text{O}\gamma 1} \rightarrow \sigma^*_{\text{N-H}}$ | 0.0219 | 60.08 |
| Thr329 A | Glu296 B | $n_{\text{O}\gamma 1} \rightarrow \sigma^*_{\text{C}\gamma\text{-H}\gamma 2}$ | 0.0007 | 2.05 |
| Val330 A | Ala293 B | $n_{\text{O}} \rightarrow \sigma^*_{\text{C}\alpha\text{-H}\alpha}$ | 0.0010 | 3.47 |
| Val330 A | Phe294 B | $n_{\text{O}} \rightarrow \sigma^*_{\text{N-H}}$ | 0.0098 | 35.56 |
| Asp256 B | Lys335 A | $n_{\text{O}\delta 1} \rightarrow \sigma^*_{\text{N}\zeta\text{-H}\zeta 1}$ | 0.0703 | 172.84 |
| Val257 B | Lys335 A | $n_{\text{O}} \rightarrow \sigma^*_{\text{C}\epsilon\text{-H}\epsilon 2}$ | 0.0012 | 4.35 |
| Val257 B | Lys335 A | $n_{\text{O}} \rightarrow \sigma^*_{\text{N}\zeta\text{-H}\zeta 2}$ | 0.0010 | 3.47 |
| Phe294 B | Thr329 A | $n_{\text{O}} \rightarrow \sigma^*_{\text{C}\gamma 2\text{-H}\gamma 22}$ | 0.0013 | 2.89 |
| Phe294 B | Val330 A | $n_{\text{O}} \rightarrow \sigma^*_{\text{N-H}}$ | 0.0090 | 30.84 |
| Phe294 B | Ser331 A | $n_{\text{O}} \rightarrow \sigma^*_{\text{C}\beta\text{-H}\beta 3}$ | 0.0020 | 6.74 |
| Gln297 B | Thr328 A | $n_{\text{O}} \rightarrow \sigma^*_{\text{N-H}}$ | 0.0169 | 59.37 |
| Gln297 B | Thr328 A | $n_{\text{O}} \rightarrow \sigma^*_{\text{O}\gamma 1\text{-H}\gamma 1}$ | 0.0011 | 3.68 |
| Thr326 B | Asn298 A | $n_{\text{O}} \rightarrow \sigma^*_{\text{C}\beta\text{-H}\beta 3}$ | 0.0020 | 6.57 |
| Thr328 B | Gln297 A | $n_{\text{O}} \rightarrow \sigma^*_{\text{N-H}}$ | 0.0114 | 40.29 |
| Thr329 B | Leu295 A | $n_{\text{O}\gamma 1} \rightarrow \sigma^*_{\text{C}\delta 1\text{-H}\delta 13}$ | 0.0006 | 1.92 |
| Thr329 B | Glu296 A | $n_{\text{O}\gamma 1} \rightarrow \sigma^*_{\text{N-H}}$ | 0.0183 | 54.52 |
| Val330 B | Ala293 A | $n_{\text{O}} \rightarrow \sigma^*_{\text{C}\alpha\text{-H}\alpha}$ | 0.0009 | 2.97 |
| Val330 B | Phe294 A | $n_{\text{O}} \rightarrow \sigma^*_{\text{N-H}}$ | 0.0099 | 36.61 |

The *n*_O_ → σ*_Cβ-Hβ3_ interactions in the Asn298 A-Thr326 B pair has *E*^(2)^ of 6.57 kJ/mol and *q*_CT_ of 0.0020 e, whereas *q*_CT_ (0.0012 *e*) and *E*^(2)^ (4.10 kJ/mol) of this interactions in the Asn298 D-Thr326 C pair are smaller. In accordance with the |*E*_HB_| prediction, the strength of this interaction in model III is thus greater than in model II. The *n*_Oγ1_ → σ*_N-H_ interaction in the Glu296 B-Thr329 A pair is the consequence of a charge transfer (0.0219 *e*) from *n*_Oγ1_ of Thr329 A to σ*_N-H_ of Glu296 B with *E*^(2)^ 60.08 kJ/mol. An attractive interaction between the interacting local orbitals of this pair is stronger than this interaction in the Glu296 A-Thr329 B pair with *q*_CT_ 0.0183 *e* and *E*^(2)^ 54.52 kJ/mol. Based on *q*_CT_ and *E*^(2)^ values, both these interactions in model III are weaker than those in model I (Table 3). Similar to model I, the *n*_Oδ1_ → σ*_Nζ-Hζ1_ interaction with *q*_CT_ 0.0703 *e* and *E*^(2)^ 172.84 kJ/mol in the Lys335 A-Asp256 B pair is the strongest donor-acceptor interaction in model III.

Our NBO results show that total stabilization energies of the charge-transfer interactions emerging from models I, II and III are 695.25, 438.75 and 651.74 kJ/mol, respectively. It is hence expected that the total donor-acceptor interactions arising from model I are stronger than those presented in models II and III.

### Comparison of stabilities of the dimeric interfaces of two conformations of PvNVPd

QTAIM and NBO analyses identified H-bonding interactions of various types affecting the stability and preservation of the dimeric interfaces within the tetrameric and the dimeric conformations of PvNVPd. To determine the strength of the intermolecular interaction, we calculated the interaction energy of each residue pair and corrected for the basis-set superposition error (BSSE) with the counterpoise (CP) correction method [28,29]. The intermolecular BSSE (IBSSE) of the interaction energy at the M06-2X/6-31G** level is defined in this equation [29]:

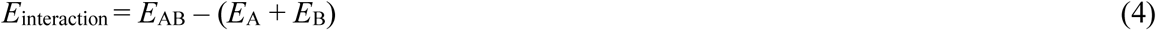

in which *E*_AB_ represents the single-point energy (SPE) of each residue pair, *E*_A_ and *E*_B_ are SPEs of the isolated residues. The moduli of interaction energies, |*E*_interaction_|, and dipole moments of residue pairs involved in the dimeric interactions in all three dimeric interfaces are tabulated in Table 5.

**Table 5.** Moduli of calculated BSSE-corrected interaction energies and dipole moments of reside pairs involved in dimeric interactions in structural models I, II, and III, assessed at the M06-2X/6-31G** level.

| Residue pair | $\vert E_{\text{Interaction}}\vert $ (kJ/mol) | | | Dipole (debye) | | |
| --- | --- | --- | --- | --- | --- | --- |
|  | Model I | Model II | Model III | Model I | Model II | Model III |
| Val257-Lys335 | 72.44 | - | - | 10.42 | - | - |
| Ala293-Val330 | 0.45 | - | 3.11 | 7.08 | - | 3.74 |
| Phe294-Thr328 | 11.79 | 13.75 | - | 3.50 | 3.64 | - |
| Phe294-Thr329 | 13.72 | 8.70 | 9.60 | 3.83 | 3.58 | 4.00 |
| Phe294-Val330 | 17.54 | 22.23 | 67.97 | 2.78 | 3.12 | 2.97 |
| Leu295-Thr329 | 20.17 | 5.69 | - | 2.28 | 2.10 | - |
| Glu296-Thr329 | 43.97 | 22.05 | 30.70 | 24.87 | 25.43 | 25.52 |
| Gln296-Thr328 | 52.47 | 48.36 | 48.69 | 6.52 | 4.17 | 2.87 |
| Asn298-Thr326 | - | 22.51 | 18.83 | - | 2.89 | 2.54 |
| Pro325-Gln300 | - | 12.92 | - | - | 5.65 | - |
| Thr326-Asn298 | - | 26.84 | - | - | 4.41 | - |
| Thr328-Gln297 | 45.58 | 40.24 | 44.69 | 6.30 | 5.31 | 3.66 |
| Thr329-Phe294 | - | 6.49 | 14.69 | - | 3.90 | 2.04 |
| Thr329-Leu295 | - | 4.78 | - | - | 1.89 | - |
| Thr329-Glu296 | 19.27 | 23.16 | 14.51 | 22.35 | 24.71 | 25.56 |
| Val330-Ala293 | 0.79 | 1.92 | 25.64 | 4.24 | 3.54 | 4.24 |
| Val330-Phe294 | - | 44.30 | 23.28 | - | 3.00 | 1.34 |
| Ser331-Phe294 | - | 8.61 | 6.80 | - | 3.29 | 1.70 |
| Lys335-Asp256 | 467.04 | - | 467.28 | 14.00 | - | 12.43 |
| Lys335-Val257 | 88.40 | - | 64.06 | 6.16 | - | 5.10 |
| Ile364-Val257 | 1.76 | 2.21 | - | 5.24 | 2.85 | - |
| Ala366-Asp256 | 16.21 | 13.24 | - | 18.64 | 18.81 | - |

Because there exists a single proton (positive charge) on the Nζ atom of Lys, Val257 A-Lys335 B and Lys335 A-Val257 B pairs in models I and III are positively charged residue pairs. In contrast, there is a single delocalized electron (negative charge) between Oε1 and Oε2 atoms of Glu; Asp possesses a delocalized negative charge between its Oδ1 and Oδ2 atoms. Glu296-Thr329, Thr329-Glu296, and Ala366-Asp256 pairs of all three models are hence negatively charged residue pairs. This charge allows the electrostatic interactions of type charge-dipole and dipole-dipole to occur in each charged residue pair. Of these, the Lys335 A-Val257 B pair of model I with |*E*_interaction_| 88.40 kJ/mol is the most stable charged residue pair in the dimeric interfaces of PvNVPd (Table 5). As a consequence, this pair plays a fundamental role in preserving the dimeric interface between subunits A and B of tetrameric PvNVPd through two moderate H-bonds Cε-Hε2…O and Nζ-Hζ2…O as well as strong charge-dipole and dipole-dipole interactions. QTAIM and NBO results prove that Nζ-Hζ2…O and Cε-Hε2…O in the Lys335 B-Val257 A and Lys335 A-Val257 B pairs of models I and III are weak H-bonds. The large values of |*E*_interaction_| in these pairs indicate that charge-dipole and dipole-dipole interactions make major contributions to the intermolecular interactions of each of these residue pairs.

In each Lys335 A-Asp256 B pair, it is possible that an electrostatic interaction of charge-charge type occurs between the localized positive charge of Lys335 A and the delocalized negative charge of Asp256 B. Large |Δ*E*_ele_| values of these pairs in model I (427.611 kJ/mol) and model III (423.14 kJ/mol) reveal that the charge-charge interactions are major contributors to the intermolecular interactions in these two pairs. In models I and III, the largest |*E*_interaction_| values pertain to the Lys335 A-Asp256 B pairs (Table 5). These pairs consequently play important roles in maintaining the dimeric interfaces between subunits A and B in both conformations of PvNVPd through charge-charge, charge-dipole, dipole-dipole, and the strong H-bond of Nζ-Hζ1…Oδ1. Identical |*E*_interaction_| values of these two pairs denote that the strengths of their intermolecular interactions are equal.

As is known, the side chains of residues Gln and Asn contain carboxamide groups; side chains of residues Tyr and Ser have hydroxyl groups. Gln296-Thr328, Asn298-Thr326, Thr326-Asn298, and Thr328-Gln297 pairs in all three models are hence polar residue pairs. Moreover, polar side chains of these residues induce a dipole moment to non-polar residues in Phe294-Thr328, Phe294-Thr329, Leu295-Thr329, Pro325-Gln300, Thr329-Phe294, Thr329-Leu295 and Ser331-Phe294 pairs. The electrostatic interactions of these residue pairs therefore include dipole-dipole and dipole-induced dipole interactions. Of these, Gln296-Thr328 pairs in all three models have the strongest intermolecular interactions because their |*E*_interaction_| values are the largest among the polar residue pairs (Table 5).

Ala293-Val330, Phe294-Val330, Val330-Ala293, Val330-Phe294, and Ile364-Val257 pairs are non-polar residue pairs in all three models. In addition to H-bonds identified, the side chains of these residues are involved in van der Waals interactions of hydrophobic type with each other. Among them, the Phe294 A-Val330 B pair of model III with |*E*_interaction_| 67.97 kJ/mol is the most stable non-polar residue pair in the dimeric interfaces of PvNVPd. The weakest intermolecular interactions exist in Ala293 A-Val330 B, Val330 A-Ala293 B, Val330 C-Ala293 D, Ile364 A-Val257 B and Ile364 C-Val257 D pairs of models I and II because of the small values of their |*E*_interaction_| (Table 5).

In the tetrameric protein, the total interaction energies pertaining to the interacting residue pairs in the dimeric A/B interface (871.59 kJ/mol) are more than twice that found in the dimeric C/D interface (328.01 kJ/mol), whereas its value in the dimeric A/B interface of the parallel model is 839.85 kJ/mol. In confirmation of the QTAIM and NBO results, the total calculated interaction energies thus emphasize that the dimeric A/B interface of tetrameric protein is the most stable interface of PvNVPd. In this interface, Val257 A-Lys335 B, Glu296 A-Thr329 B, Gln296 A-Thr328 B, Thr328 A-Gln297 B, Lys335 A-Asp256 B and Lys335 A-Val257 B pairs are crucial residue pairs playing significant roles in the stabilization of the dimeric A/B interface through van der Waals, charge-charge, charge-dipole, dipole-dipole, and H-bonding interactions. Although the calculated interaction energies establish that the dimeric interface between subunits C and D is the weakest interface of PvNVPd, the dimeric interactions resulting from the residue pairs of both dimeric interfaces provide the formation and conservation of quaternary structure of tetrameric protein.

In parallel model, Phe294 A-Val330 B, Glu296 A-Thr329 B, Gln296 A-Thr328 B, Thr328 A-Gln297 B, Val330 A-Ala293 B, Lys335 A-Asp256 B and Lys335 A-Val257 B pairs are key residue pairs. Charge-charge, charge-dipole, dipole-dipole and H-bonding interactions derived from these residue pairs ensure the stability and preservation of the parallel dimeric A/B interface.

## Theoretical methods

As the positions of hydrogen atoms were not determined in the mentioned crystal structures from X-ray diffraction, the hydrogen atoms of each structural model were optimized using a hybrid meta–GGA density functional (M06-2X) [18,19] in conjunction with a standard 6-31G** basis set to ensure that their positions were reasonable. During the geometry optimization, the positions of all other atoms remained frozen. The vibrational frequencies were also calculated for the optimized structural model at the respective level; all stationary points were found to be true minima (number of imaginary frequencies was zero). QTAIM and NBO analyses were performed on each wave function derived from each optimized structural model at the respective level.

All DFT calculations were implemented with the Gaussian 09 program package [30]; QTAIM calculations were made with AIM 2000 software [31].

## Conclusion

We have applied QTAIM and NBO analyses to identify precisely the dimeric interactions of various types that stabilize dimeric interfaces in the dimeric and tetrameric conformations of PvNVPd. QTAIM and NBO analyses indicated that all three dimeric interfaces have H-bonds of common types with strength ranging from weak to strong. In the dimeric A/B interfaces of both conformations, Lys335-Asp256 and Lys335-Val257 pairs are two important residue pairs playing significant roles in maintaining these interfaces especially through strong charge-charge and charge-dipole interactions, as well as a strong H-bond of Nζ-Hζ1…Oδ1.

Moreover, Phe294-Val330, Gln296-Thr328, Glu296-Thr329, Thr328-Gln297, and Val330-Ala293 pairs are critical residue pairs in all three dimeric interfaces, providing their stabilities through hydrophobic, charge-dipole, dipole-dipole and H-bonding interactions. Side chains of residues Val257, Ala293, Phe294, Val330 and Ile364 contribute to the hydrophobic nature of each of these dimeric interfaces.

The highest strengths of the intermolecular dimer–dimer interactions are identified in the dimeric interface between subunits A and B of the tetrameric conformation of PvNVPd. The dimeric A/B interface of the tetrameric protein is consequently the most stable interface of PvNVPd.

## Acknowledgment

We are indebted to the supporting staffs at beamlines TPS 05A, BL13B1 and BL15A1 at the National Synchrotron Radiation Research Center (NSRRC) and Masato Yoshimura at the beamlines BL12B2 and BL44XU for the structure determination of PvNV and the P-domains. We thank Saeid MalekZadeh for the assistance during manuscript preparation and calculation. We also thank the support of computation facilities at NSRRC. This work was supported in part by Ministry of Science and Technology (MOST) grants 105-2311-B-213-001-MY3, 107-2923-B-213-001-MY3 and 108-2311-B-213-001-MY3, and NSRRC grants to C.-J. Chen.

